# Enhancing plant broad-spectrum resistance through engineered pattern recognition receptors

**DOI:** 10.1101/2025.02.26.640353

**Authors:** Yuankun Yang, Christina E. Steidele, Birgit Löffelhardt, Lili Huang, Jonathan D.G. Jones, Thorsten Nürnberger, Pingtao Ding, Andrea A. Gust

**Affiliations:** Department of Plant Biochemistry, Center of Plant Molecular Biology (ZMBP), Eberhard-Karls-University of Tübingen, Tübingen, Germany; Institute of Biology Leiden, Leiden University, Leiden, The Netherlands; Chair of Phytopathology, TUM School of Life Sciences, Technische Universität München, Freising-Weihenstephan, Germany; State Key Laboratory of Crop Stress Biology for Arid Areas, College of Plant Protection, Northwest A&F University, Yangling, Shaanxi, China; The Sainsbury Laboratory, University of East Anglia, Norwich, UK

## Abstract

Conventional plant resistance breeding has primarily focused on intracellular immune receptors, while cell-surface pattern recognition receptors (PRRs) have been underexplored due to their comparatively modest contributions to resistance. However, PRRs offer significant untapped potential for crop improvement. In this study, we demonstrate that the Arabidopsis receptor-like protein RLP23, which recognizes molecular patterns from three distinct microbial kingdoms, confers broad-spectrum resistance when introduced into the Solanaceae crop tomato. We also identify the intracellular (IC) domain of RLP23 as crucial for ensuring compatibility and efficacy during heterologous expression. Targeted engineering of the IC domain significantly enhances RLP23 potential utility, enabling transfer of robust resistance against bacterial, fungal, and oomycete pathogens to other plants without compromising yield. We extended this RLP engineering strategy to rice and poplar, highlighting its broad applicability. These findings establish a versatile framework for PRR-based engineering, opening new avenues for sustainable crop protection.

## Introduction

Plants rely on a limited set of immune receptors to combat diverse pathogens, classified into pattern recognition receptors (PRRs), which mediate cell-surface immunity (pattern-triggered immunity, PTI), and nucleotide-binding leucine-rich repeat receptors (NLRs), which control intracellular immunity (effector-triggered immunity, ETI)^1^. Evolutionary divergence among plant species results in varying immune receptor distributions and resistance traits, making interspecies receptor transfer a promising strategy to boost crop resilience.

Over the past decades, many resistance genes, mainly from the NLR family, have been cloned and used in crop breeding. NLRs typically confer race-specific resistance by recognizing specific pathogen effectors, but the rapid evolution of pathogens can render NLR-equipped plants vulnerable to newly emerging pathogen strains^2^. To address this, stacking multiple NLR genes has been explored as a strategy to enhance resistance^3^, though the continuous evolution of pathogens poses a challenge to the durability of such crops.

In contrast, PRRs recognize conserved pathogen-associated molecular patterns (PAMPs), which are crucial to microbial physiology and less prone to evolution, making them attractive for obtaining broad-spectrum resistance. Despite this advantage, PRRs have been underutilized in crop breeding due to their relatively modest immune responses compared to intracellular receptors. Only a few PRRs, such as the Arabidopsis receptor-like kinase (RLK) EF-TU RECEPTOR (EFR), have been used to enhance bacterial disease when expressed in crops like wheat, apple, and tomato^4,5^.

Leucine-rich repeat receptor-like proteins (LRR-RLPs) also show potential for improving disease resistance. For example, transferring the elicitin receptor ELICITIN RESPONSE (ELR) from wild potato to cultivated varieties confers resistance to *Phytophthora infestans*^6^. Similarly, introducing the xyloglucanase receptor RESPONSE TO XEG1 (RXEG1) from *Nicotiana benthamiana* into wheat enhances resistance to *Fusarium* head blight^7^. However, these PRRs are pathogen-specific, limiting their broader application^6,7^.

Two Arabidopsis receptors, RLP23 and RLP30, detect molecular patterns from all three microbial kingdoms^8,9^. RLP23 senses Necrosis- and ethylene-inducing peptide 1-like proteins (NLPs) from bacteria, fungi, and oomycetes, while RLP30 recognizes small cysteine-rich proteins (SCPs) from fungal and oomycete pathogens, along with an unidentified *Pseudomonas* pattern. This broad recognition makes RLP23 and RLP30 promising candidates for PRR-based crop engineering. Notably, transgenic potato expressing RLP23 shows elevated resistance to the fungal pathogen *Sclerotinia sclerotiorum* and the oomycete *Phytophthora infestans*^8^.

However, heterologous expression of RLPs often faces compatibility issues, requiring co-expression of additional receptor complex components for optimal functionality. For instance, co-expressing RLP30 with the adapter kinase SUPPRESSOR OF BIR1 1 (SOBIR1) from *Arabidopsis thaliana* in *N. tabacum* achieved stronger resistance, compared to RLP30 alone^9^. Yet, overexpression of co-receptors can result in autoimmune phenotypes, such as dwarfism and premature flowering^10^.

While recent advances in NLR engineering have expanded their recognition spectrum, as demonstrated for the rice NLR Pik-1^11^, they remain pathogen-specific. In contrast, engineering PRRs for broad-spectrum recognition while enhancing their functionality is an underexplored approach for crop breeding.

Here, we show that the intracellular (IC) domain of RLPs determines compatibility during heterologous expression. By modifying the IC domain of RLP23, we significantly enhanced tomato resistance to bacteria, fungi, and oomycetes, without compromising yield. Notably, this PRR modification strategy is also applicable to the monocot species rice and the perennial poplar. Our findings provide a scalable framework for engineering PRRs to achieve broad-spectrum resistance, offering a sustainable approach for crop improvement.

## Results

### RLP23 transformation enhances tomato resistance to multiple pathogens

The Arabidopsis receptor RLP23 recognizes the conserved immunogenic epitope nlp20, which is present in pathogenic bacteria, fungi, and oomycetes^12^, making it a promising tool for crop improvement. To explore this potential, we generated a transgenic tomato line stably expressing RLP23 fused with GFP under the constitutive 35S promoter (*p35S::RLP23-GFP*). Western Blot analyses confirmed successful expression of RLP23 (Supplementary Fig. 1a, b).

Compared to wild-type tomato plants, the transgenic line showed increased ethylene production in response to nlp20 stimulation (Fig. 1a). To assess whether RLP23-mediated recognition of nlp20 enhances immunity, we challenged the transgenic plants with the bacterial pathogen *Pseudomonas syringae* pv. *tomato* (*Pst*) DC3000, the fungal pathogen *Botrytis cinerea*, and the oomycete *Phytophthora infestans*. Notably, transgenic tomatoes displayed enhanced antibacterial immunity against *Pst* DC3000. Additionally, RLP23-expressing plants showed significantly smaller disease lesions upon infection with both *B. cinerea* and *P. infestans* compared to wild-type plants (Fig. 1b-d), confirming that RLP23 expression confers broad-spectrum resistance to diverse pathogens in tomato.

**Fig. 1:**
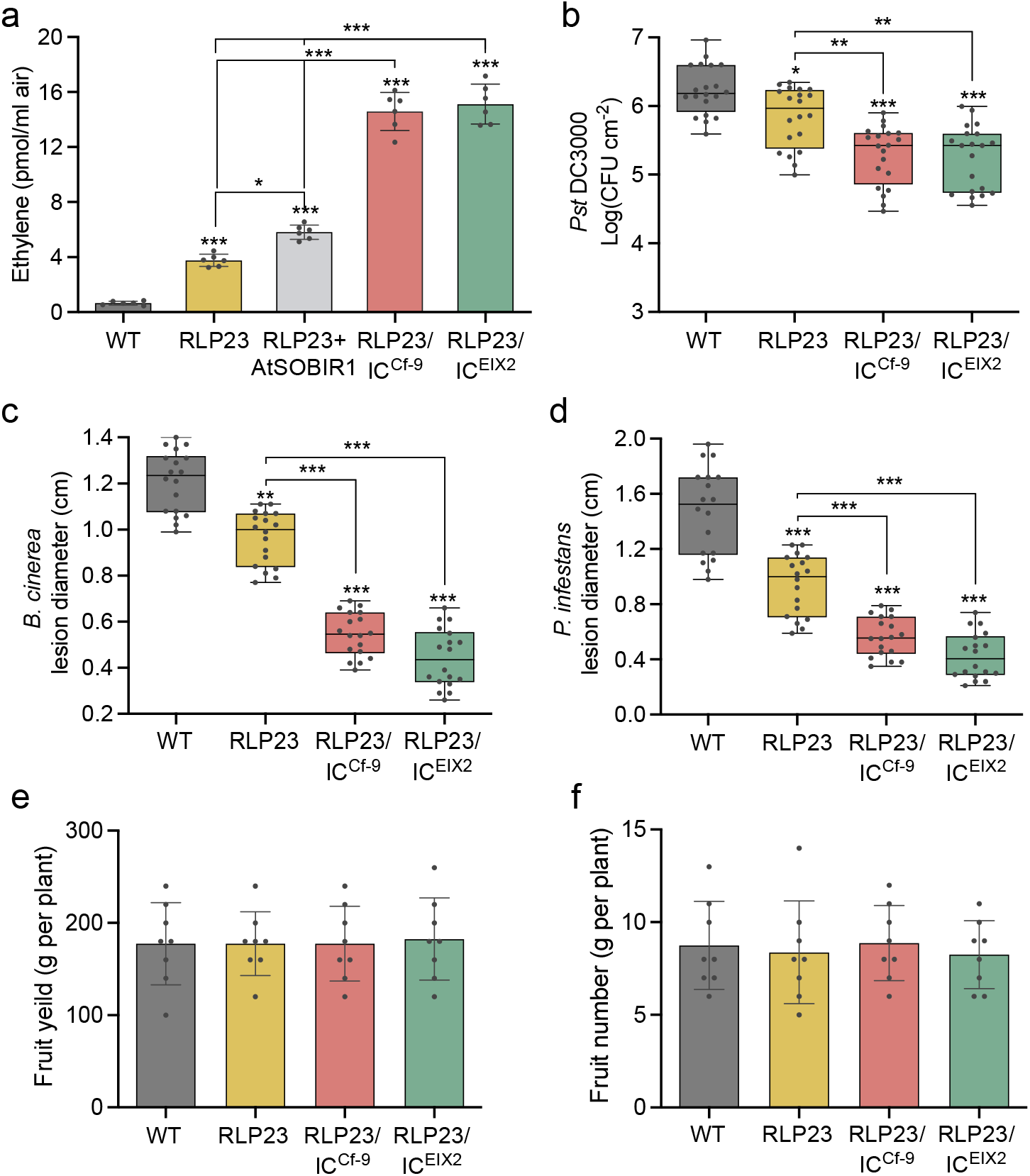
Expression of RLP23 chimeras in tomato confers increased defense without compromising yield. **a**, Ethylene accumulation in wild-type (WT) and transgenic tomato plants expressing RLP23, RLP23 with AtSOBIR1, RLP23/IC^Cf-9^, or RLP23/IC^EIX2^, measured after 4 hours of treatment with 2 µM nlp20. **b**, Bacterial growth of *Pseudomonas syringae* pv. *tomato* (*Pst*) DC3000 in wild-type (WT) and transgenic tomato plants, quantified as colony-forming units (CFU) in leaf extracts three days post-inoculation. **c**,**d**, Lesion diameters on wild-type (WT) and transgenic tomato leaves infected with *Botrytis cinerea* (c) or *Phytophthora infestans* (d), assessed two days after drop inoculation. **e**,**f**, Total weight (e) and number (f) of mature fruits harvested per tomato plant. Data points are represented as dots (*n* = 6 for a; *n* = 20 for b; *n* = 18 for c, d; *n* = 8 for e, f). For panels b–d, data are displayed as box plots (center line, median; bounds of box, the first and third quartiles; whiskers, 1.5 times the interquartile range; error bar, minima and maxima). Statistically significant differences from WT are indicated (*P ≤ 0.05, **P ≤ 0.01, ***P ≤ 0.001; Tukey’s multiple comparisons test). Each experiment was repeated three times with consistent results.

To further explore whether co-expression of RLP23 with the adaptor protein AtSOBIR1 could boost immunity, we generated a transgenic tomato line expressing both RLP23 and AtSOBIR1. Co-expression resulted in a modest increase in nlp20-induced ethylene production compared to RLP23 expression alone (Fig. 1a). However, AtSOBIR1 expression also caused dwarfism, likely due to autoimmune activation (Supplementary Fig. 1a). This phenotype contrasts with *N. tabacum*, where co-expression of RLP30 and AtSOBIR1 did not affect plant growth^9^, suggesting species-specific differences in AtSOBIR1-mediated immune regulation. While co-expression with AtSOBIR1 slightly enhanced immune signaling, its negative impact on growth limits its practical application for crop improvement. Overall, these results demonstrate that stable ectopic expression of RLP23 in tomato provides broad spectrum resistance to destructive bacterial, fungal, and oomycete pathogens.

### Full RLP functionality in heterologous plants depends on the IC domain

LRR-RLPs contain a short C-terminal intracellular (IC) domain (Supplementary Fig. 2a), but its role in immune signaling remains unclear. To investigate its contribution to RLP-mediated immunity, we introduced a truncated version of RLP23 lacking the IC domain (RLP23ΔIC) into the Arabidopsis *rlp23*-1 mutant. However, deletion of the IC domain did not impair ethylene accumulation in response to nlp20 treatment (Supplementary Fig. 2b). Surprisingly, transient expression of RLP23ΔIC in *N. benthamiana* resulted in a significantly weaker ethylene response to nlp20 compared to full-length RLP23 (Supplementary Fig. 2c,f), suggesting that the IC domain plays a species-dependent role in immune activation.

To further explore this, we examined RLP30, an SCP receptor with a short IC domain of 23 amino acids^9^. In *N. benthamiana* plants lacking the SCP receptor RE02 (ΔRE02), expression of full-length RLP30 successfully restored SCP-triggered ethylene production (Supplementary Fig. 2d,e,g,h). However, expression of RLP30 lacking the IC domain (RLP30ΔIC) in ΔRE02 plants partially reduced the SCP-induced immune response (Supplementary Fig. 2d,g). These findings suggest that while the IC domain plays a supportive role, it is not essential for RLP-mediated immunity in heterologous plant systems.

Interestingly, the RLP30 IC domain contains four putative phosphorylation sites (T761, T783, T784, S785). To determine if phosphorylation regulates immune activation, we generated an alanine-substituted phospho-null mutant at these sites (4mut-A). However, SCP-triggered immune responses remained unchanged in the mutant (Supplementary Fig. 2e,h), indicating that IC domain-mediated immunity operates independently of phosphorylation events.

### IC domain swaps modulate RLP functionality

The reduced activity of RLP23ΔIC and RLP30ΔIC in *N. benthamiana* underscores the critical role of the IC domain in heterologous expression systems. To assess whether IC domains from solanaceous RLPs could enhance RLP23 functionality in the Solanaceae family, we selected two well-characterized tomato RLPs, EIX2 and Cf-9, as IC domain donors due to their structural similarity to RLP23. EIX2 detects ethylene-inducing xylanase (EIX), while Cf-9 recognizes the *Cladosporium fulvum* effector protein Avr9^13,14^.

To investigate the impact of these IC domains, we constructed chimeric receptors by swapping the IC domains of RLP23, EIX2, and Cf-9. The chimeras, RLP23/IC^EIX2^ and RLP23/IC^Cf-9^, fused RLP23 with the IC domains of EIX2 or Cf-9, while EIX2/IC^RLP23^ and Cf-9/IC^RLP23^ carried the RLP23 IC domain in place of their native C-termini (Fig. 2a). Expression of RLP23/IC^EIX2^ and RLP23/IC^Cf-9^ in *N. benthamiana* resulted in a fourfold increase in ethylene production in response to nlp20 treatment compared to wild-type RLP23 (Fig. 2b), indicating that the IC domains from solanaceous RLPs enhance RLP23-mediated immune responses. In contrast, replacing the IC domain in EIX2 and Cf-9 with that of RLP23 significantly reduced their respective responses to EIX and Avr9 (Fig. 2c,d), demonstrating that the IC domain contributes to receptor-specific immune signaling.

**Figure 2:**
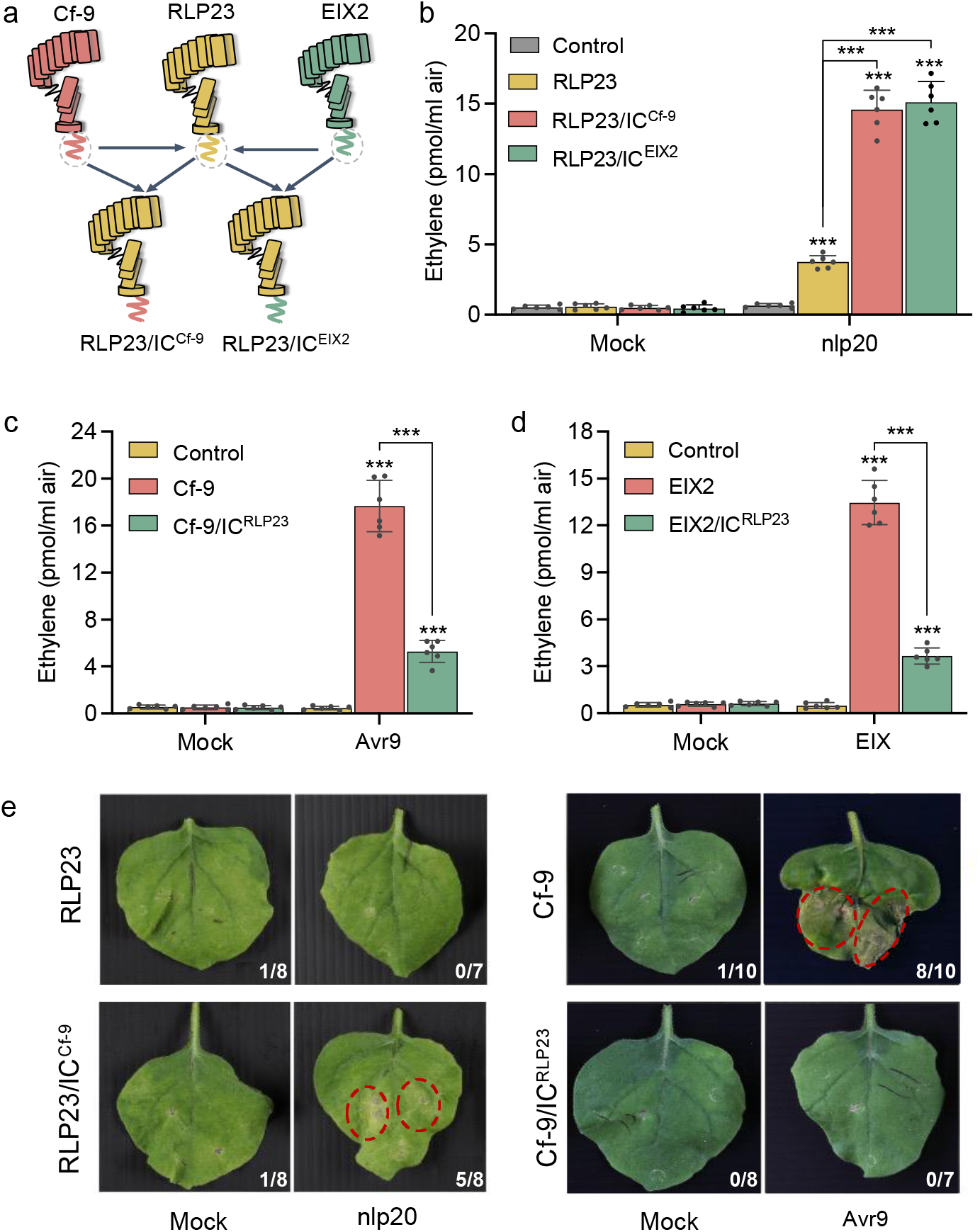
IC domain swaps result in RLP activity differences in *N. benthamiana*. **a**, Schematic representation of RLP23, Cf-9, and EIX2, along with their chimeric variants featuring reciprocal intracellular domain swaps. **b**, Ethylene accumulation in *N. benthamiana* leaves transiently expressing GFP (Control), RLP23-GFP, RLP23/IC^Cf-9^-GFP, or RLP23/IC^EIX2^-GFP, measured 4 hours after treatment with water (mock) or 2 µM nlp20. **c**, Ethylene accumulation in *N. benthamiana* leaves expressing GFP (Control), Cf-9-GFP, or Cf-9/IC^RLP23^-GFP, assessed 4 hours after treatment with 5 µl of *N. tabacum* apoplastic fluid with or without Avr9. **d**, Ethylene accumulation in *N. benthamiana* leaves expressing GFP (Control) or indicated RLP-GFP constructs, measured 4 hours after treatment with water (mock) or 1 µM EIX. **e**, Cell death development in *N. benthamiana* leaves transiently expressing indicated RLPs, 24 hours after infiltration with water (mock), 20 µM nlp20, or 100 µl apoplastic fluid with or without Avr9. The number of necrotic leaves is presented relative to the total number of infiltrated leaves. Data points are represented as dots (*n* = 6 for b–d). Statistically significant differences from control plants are indicated (Tukey’s multiple comparisons test, ***P ≤ 0.001). Shown is one out of three experiments with similar results.

Interestingly, RLP23/IC^Cf-9^ expression induced nlp20-triggered cell death, a hallmark of ETI which has not been previously observed in any plant following nlp20 treatment (Fig. 2e). In contrast, Cf-9/IC^RLP23^ did not produce the cell death response usually triggered by Avr9 (Fig. 2e), indicating that the IC domain plays a key role in regulating immune signaling outcomes. These findings highlight the functional importance of the IC domain in modulating RLP activity, enhancing immune responses, and influencing cell death phenotypes in heterologous expression systems.

### Disease resistance in tomato enhanced by RLP23 IC domain editing without yield loss

Building on the enhanced RLP23 functionality observed in *N. benthamiana*, we explored whether RLP23 chimeric receptors could improve disease resistance in tomato. The RLP23/IC^EIX2^ and RLP23/IC^Cf-9^ constructs were stably introduced into the tomato cultivar Moneymaker, which does not naturally respond to nlp20. Consistent with the results from *N. benthamiana*, transgenic tomato lines expressing the RLP23 hybrid receptors showed a substantial increase in nlp20-induced ethylene production and enhanced disease resistance (Fig. 1a-d). Compared to wild-type and RLP23-expressing plants, RLP23/IC^EIX2^ and RLP23/IC^Cf-9^ transgenic tomatoes exhibited significantly reduced bacterial growth after *Pst* DC3000 infection (Fig. 1b) and developed smaller disease lesions when challenged with the necrotrophic fungus *B. cinerea* (Fig. 1c) or the oomycete *P. infestans* (Fig. 1d).

The trade-off between defense and growth has long been a key challenge in crop breeding^15^. In contrast to the dwarfism observed in plants co-expressing RLP23 with AtSOBIR1, tomato lines carrying RLP23/IC^EIX2^ and RLP23/IC^Cf-9^ showed no developmental defects, with growth patterns comparable to those of wild-type and RLP23-expressing plants (Supplementary Fig. 1a,b). Importantly, the enhanced resistance conferred by the engineered RLP23 receptors did not affect fruit yield. Both transgenic and wild-type tomato plants produced similar numbers and total weights of mature fruits (Fig. 1e,f). These findings demonstrate that RLP engineering can significantly enhance broad-spectrum resistance in tomato without compromising yield.

### Interaction of RLP23 with SOBIR1 is modulated by the IC domain

To understand how tomato RLP-IC domains enhance RLP23 functionality in Solanaceae, we examined their impact on immune signaling. Consistent with the elevated nlp20-triggered ethylene levels observed in RLP23-expressing plants, we also detected significant upregulation of immune-related gene transcription upon nlp20 treatment. Genes such as *PR1a* and *RIPK* were strongly induced, with even higher expression levels in transgenic tomato lines expressing the chimeric RLP23/IC^EIX2^ (Supplementary Fig. 3). Notably, these lines also showed nlp20-induced transcript accumulation of genes typically associated with ETI, such as *EDS1, NRC2*, and *NRC4a* (Supplementary Fig. 3). These findings suggest that the chimeric RLP23 variants may activate both PTI and ETI, resulting in a more robust, broad-spectrum immune response.

To assess whether ligand-binding capacities were altered in the RLP23 chimeric proteins, we tested their interaction with biotinylated nlp24 (nlp24-bio), a functional analog of nlp20 that includes a C-terminal four-amino-acid extension. All tested variants, including wild-type RLP23, RLP23/IC^EIX2^, and RLP23/IC^Cf-9^, associated similarly with nlp24-bio. Excess of untagged nlp24 effectively displaced nlp24-bio from all RLP23 variants (Fig. 3a), indicating that the IC domain replacement does not affect the ligand binding properties of RLP23. Thus, the enhanced immune response is unlikely to be due to altered ligand recognition.

**Figure 3:**
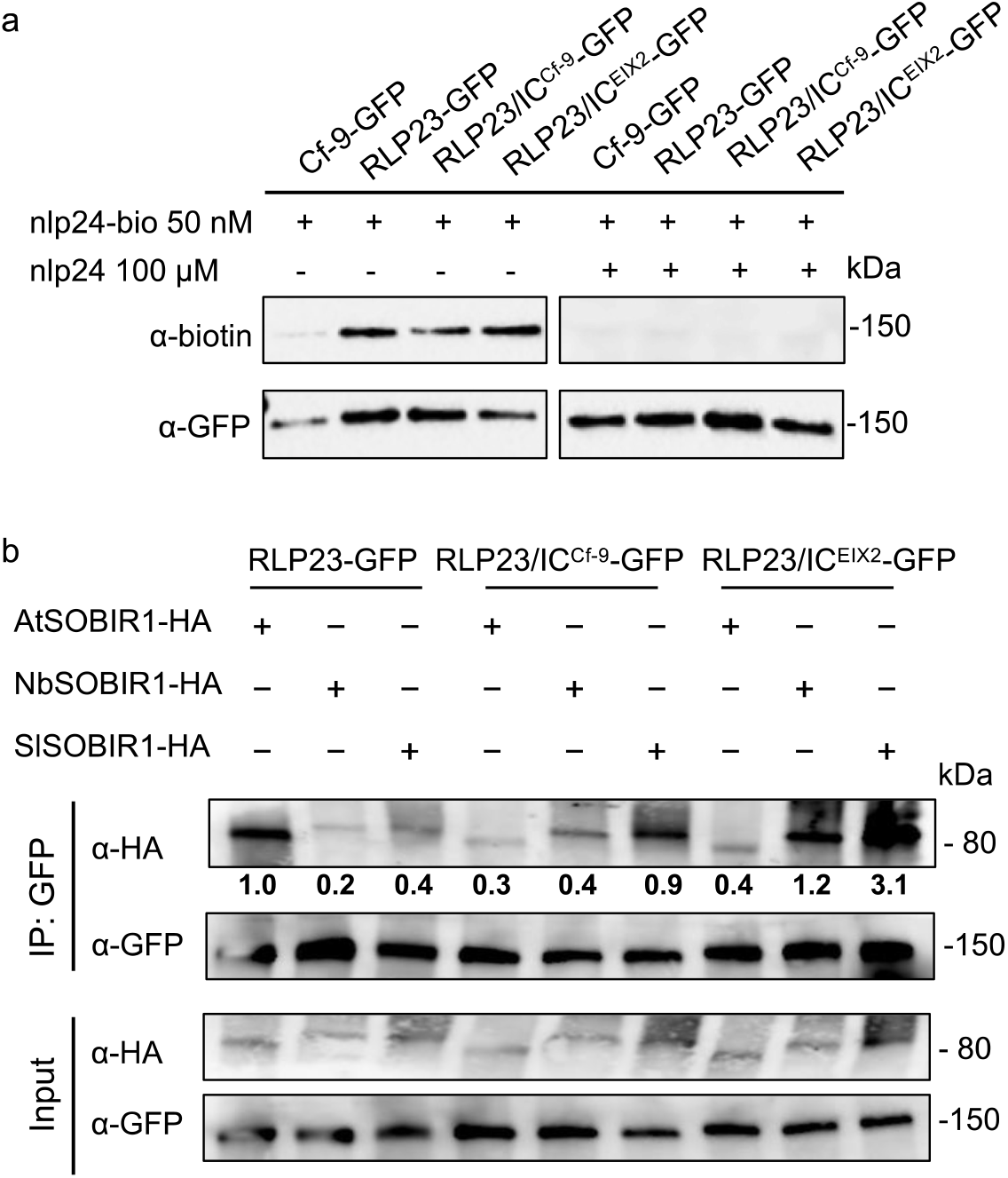
Replacement of RLP23-IC doesn’t affect ligand binding but the interaction with SOBIR1. **a**, Ligand-binding assay in *N. benthamiana* leaves transiently expressing the indicated GFP-tagged RLP variants. Leaves were treated with biotinylated nlp24 (nlp24-bio) in the presence (+) or absence (-) of unlabeled nlp24, followed by co-immunoprecipitation using a GFP-trap. **b**, Co-immunoprecipitation assay in *N. benthamiana* leaves co-expressing GFP-tagged RLP23 variants with AtSOBIR1-HA, NbSOBIR1-HA, or SlSOBIR1-HA. Leaf protein extracts (Input) were subjected to co-immunoprecipitation with GFP beads (IP:GFP) and immunoblotting using tag-specific antibodies. Protein levels were quantified relative to AtSOBIR1-HA precipitated by RLP23-GFP, set as 1. Representative results from three independent experiments are shown.

Interestingly, when we assessed the interaction between RLP23 and the adaptor kinase SOBIR1 from different plant species, we observed a stronger association of RLP23/IC^EIX2^ and RLP23/IC^Cf-9^ with tobacco SOBIR1 (NbSOBIR1) and tomato SOBIR1 (SlSOBIR1) compared to AtSOBIR1 (Fig. 3b). While wild-type RLP23 preferentially interacted with AtSOBIR1, the chimeric receptors, RLP23/IC^EIX2^ and RLP23/IC^Cf-9^, exhibited the strongest association with SlSOBIR1 (Fig. 3b), indicating that the IC domain plays a crucial role in modulating RLP-SOBIR1 interaction. In summary, engineering the IC domain can enhance RLP23’s adaptation and integration into the immune signaling networks of heterologous plants.

### RLP23 expression in rice confers recognition of NLPs from blast fungi

Given that Arabidopsis RLP23 conferred resistance in dicot plants such as tomato and tobacco (Figs. 1, 2), we next investigated its potential to improve immunity in monocot crops. The rice blast fungus *Magnaporthe oryzae* encodes the NLP protein MoNLP1, which contains the conserved immune epitope nlp20^16^. However, rice does not naturally recognize this protein (Fig. 4a). To determine whether RLP23 can mediate MoNLP1 recognition in rice, we transiently expressed the *p35S::RLP23-GFP* construct in rice protoplasts. Upon MoNLP1 treatment, RLP23 expression triggered a significant accumulation of reactive oxygen species (ROS), with no further enhancement upon co-expression with AtSOBIR1 (Fig. 4a). Moreover, MoNLP1-treated RLP23-expressing rice protoplasts showed significant upregulation of defense-related genes, including *OsWRKY70, OsPAL1, OsPR1a* and *OsMAPK6* (Fig. 4b). These results indicate that RLP23 enables MoNLP1 recognition and activates defense responses in rice.

**Figure 4:**
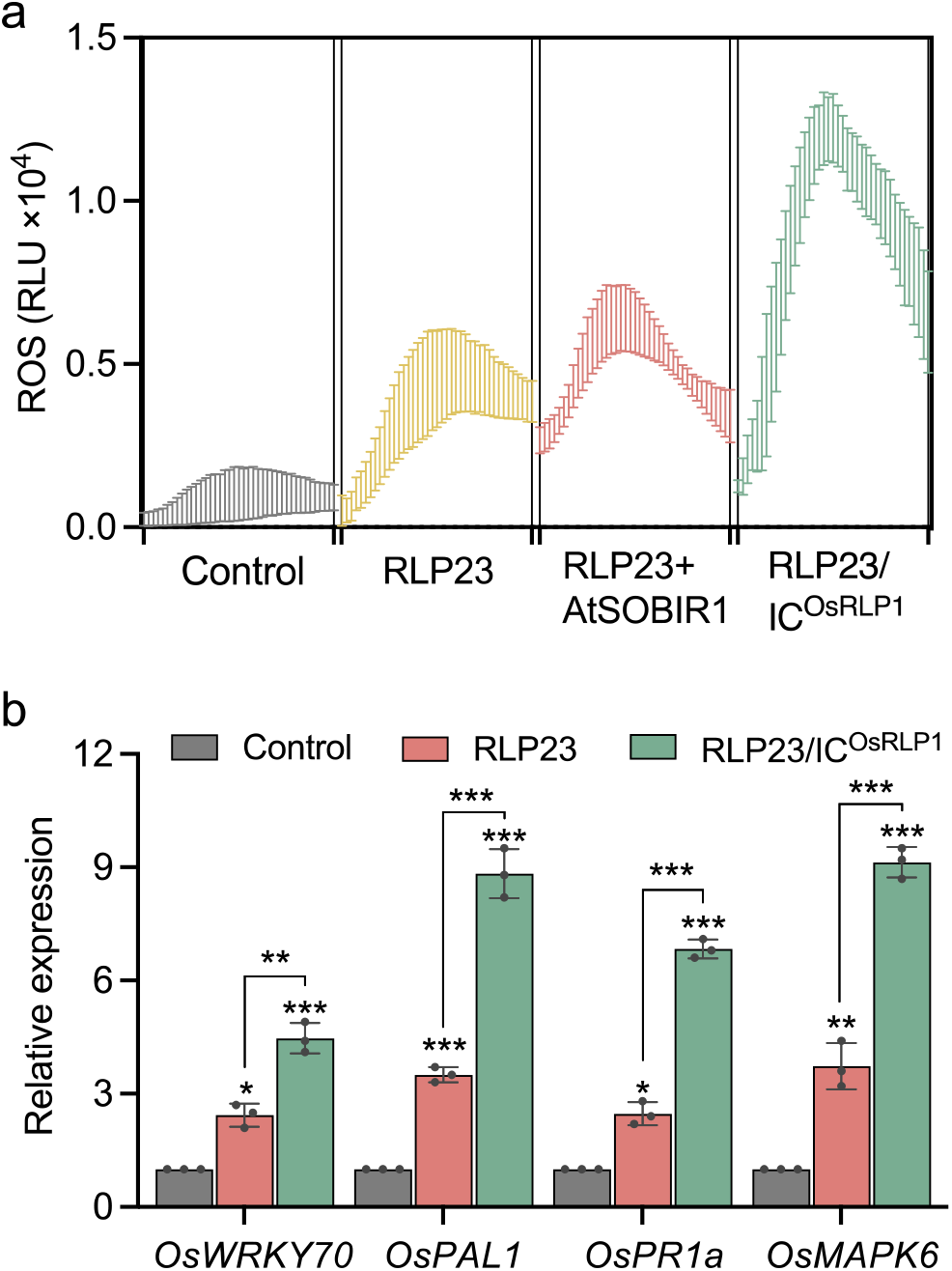
Expression of RLP23 chimeras enhances immune responses in rice. **a**, Reactive oxygen species (ROS) production in rice protoplasts expressing GFP (Control), RLP23-GFP, RLP23-GFP co-expressed with AtSOBIR1-HA, or RLP23/IC^OsRLP1^-GFP, following treatment with 5 µM purified MoNLP1. Data are presented as relative light units (RLU) over 60 minutes. **b**, RT-qPCR analysis of rice protoplasts expressing GFP (Control), RLP23-GFP, or RLP23/IC^OsRLP1^-GFP, treated with 2 µM MoNLP1 for 24 hours. Expression levels of the indicated marker genes were quantified using gene-specific primers, normalized to *OsActin* transcript levels, and presented relative to the control (set to 1). Data points are shown as dots (*n* = 3 for b). Statistically significant differences from the control are indicated (Tukey’s multiple comparisons test, *P ≤ 0.05, **P ≤ 0.01, ***P ≤ 0.001). Each experiment was performed three times with similar results.

To further improve RLP23 functionality in rice, we designed the hybrid receptor RLP23/IC^OsRLP1^ by replacing the IC domain of RLP23 with that of rice OsRLP1, a key component of antiviral immunity in rice^17^. Compared to wild-type RLP23, expression of RLP23/IC^OsRLP1^ in rice protoplasts led to further increased ROS production and stronger defense gene induction upon MoNLP1 treatment (Fig. 4a,b). These findings, consistent with our results in tomato, highlight the potential of RLP engineering to enhance disease resistance in monocot crops like rice.

### Heterologous expression of Arabidopsis RLP23 enhances fungal resistance in poplar

RLP23 is specific to the Brassicaceae family, while NLP proteins are widely conserved among fungal pathogens, including *Marssonina brunnea*, the causal agent of black spot disease in the economically important tree species poplar^18^. To explore the potential of cross-species RLP transfer for tree improvement, we cloned the NLP-type protein MbNLP1 from *M. brunnea*, which is recognized by RLP23 but not by poplar (Fig. 5a)^18^. MbNLP1 contains a conserved 24-amino acid immunogenic fragment (nlp24^MbNLP1^), which induces ROS accumulation and upregulates defense gene expression in poplar leaves transiently expressing RLP23 (Fig. 5a,b). Consistent with our results in rice, co-expression of AtSOBIR1 did not further enhance nlp24^MbNLP1^-triggered ROS production in poplar (Fig. 5a). Importantly, ectopic expression of RLP23 conferred fungal resistance, as evidenced by reduced disease symptoms in poplar leaves compared to GFP-expressing controls (Fig. 5c).

**Figure 5:**
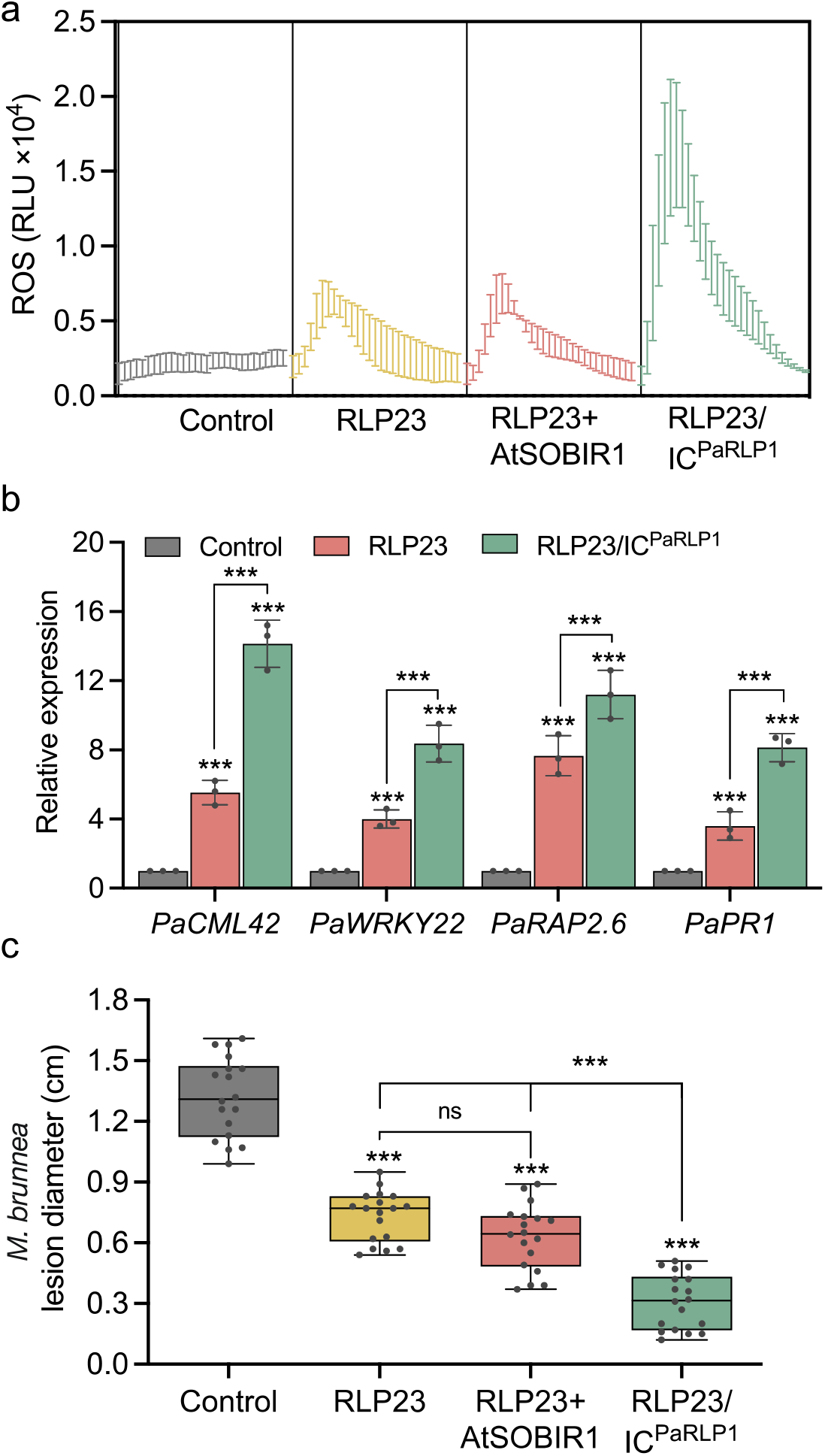
RLP23 and its hybrid form confer increased defense in poplar. **a**, ROS production in poplar leaves transiently expressing GFP (Control) or the indicated constructs for 48 hours. ROS accumulation was measured for 60 minutes following treatment with 2 µM nlp24^MbNLP1^ and is given in relative light units (RLU). **b**, RT-qPCR analysis of poplar leaves expressing GFP (Control), RLP23, or RLP23/IC^PaRLP1^ for 48 hours, followed by treatment with 1 µM nlp24^MbNLP1^ for 24 hours. Expression levels of the indicated genes were determined using gene-specific primers, normalized to *PaEF1α* transcript levels, and presented relative to the control (set to 1). **c**, Lesion diameter measured three days post-inoculation with *Marssonina brunnea* in poplar leaves transiently expressing GFP (Control) or the indicated RLP-GFP fusion constructs. Data points are shown as dots (*n* = 3 for b; *n* = 18 for c), and box plots in panel c show center line, median; bounds of box, the first and third quartiles; whiskers, 1.5 times the interquartile range; error bar, minima and maxima. Statistically significant differences from control plants are indicated (Tukey’s multiple comparisons test, ***P ≤ 0.001, ns: not significant). Shown is each one representative experiment out of three with similar results.

To further enhance RLP23 function in poplar, we engineered the chimeric receptor RLP23/IC^PaRLP1^, by fusing the intracellular domain of poplar RLP1 with RLP23^19^. Poplar leaves expressing this hybrid receptor showed higher ROS accumulation and more pronounced defense gene expression in response to nlp24^MbNLP1^ (Fig. 5a,b). These elevated immune responses translated into significantly enhanced antifungal resistance (Fig. 5c), demonstrating that RLP engineering can be effectively applied to tree breeding, offering a promising strategy for improving disease resistance in perennial crops.

## Discussion

Most resistance genes used in breeding programs originate from intracellular NLR receptors, but pathogens rapidly evolve to evade NLR-mediated detection, thus undermining resistance^20^. In contrast, PRRs detect highly conserved molecular patterns, offering more durable, broad-spectrum defense. This makes PRR-based engineering a promising strategy for sustainable crop improvement.

Among PRRs, RLP23 and RLP30 stand out for their broad-spectrum pathogen recognition across bacterial, fungal, and oomycete pathogens^8,9^. Notably, RLP23 retains functionality when transferred between species, conferring robust resistance in tomato against various pathogen classes. This aligns with previous findings in potato^8^, reenforcing RLP23 as a strong candidate for Solanaceae crop improvement. Remarkably, RLP23 also functions effectively in monocots like rice and woody plants like poplar, enabling recognition of NLP proteins from pathogens such as *M. oryzae* (rice blast) and *M. brunnea* (poplar black spot disease). The ability of RLP23-expressing poplar leaves to resist *M. brunnea* infection further supports its cross-species functionality^18^.

The intracellular (IC) domain of RLPs, though short with 10–50 amino acids^21,22^, plays a key role in heterologous expression systems. While dispensable for RLP23-mediated immunity in Arabidopsis, IC-deletion weakens RLP23 function in *N. benthamiana*, emphasizing its role in facilitating immune signalling. Chimeric RLP23 receptors with IC domains from tomato RLPs enhance immune signalling in both *N. benthamiana* and tomato, a strategy that also proved effective in rice and poplar. Similarly, Arabidopsis RLP1 becomes functional in tobacco only when fused to the C-terminus of Solanaceous RLPs like EIX2^23^.

However, RLP transfer across species is not always successful. For example, tomato Cf-9 does not confer Avr9 perception in Arabidopsis (Supplementary Fig. 4), likely due to incompatibility with Arabidopsis co-receptors. Notably, native SlSOBIR1 is essential for Cf-4 stabilization^24^. Similarly, replacing the Cf-9 IC domain with that of RLP23 or RLP30 did not restore its function (Supplementary Fig. 4), suggesting that additional species-specific factors, such as glycosylation patterns, may affect RLP folding and stability. Moreover, the Cf-9 IC domain disrupts RLP23 function in Arabidopsis, highlighting the complexities of cross-species RLP adaptation.

Structural analyses show that the IC domains of tomato EIX2 (37 aa) and Cf-9 (30 aa) are longer than that of RLP23 (19 aa), which may enhance interactions with Solanaceae SOBIR1. Co-immunoprecipitation assays confirm that IC domain modifications improve RLP-SOBIR1 binding without altering ligand recognition, which remains determined by the extracellular domain^22^.

Importantly, engineered “super” tomato lines exhibit broad-spectrum immunity to bacterial, fungal, and oomycete pathogens without compromising growth or yield – a major milestone in disease resistance breeding. Furthermore, RLP23 activation appears to synergize both cell-surface and intracellular immunity (PTI and ETI), amplifying defense outputs^25^. Elevated expression of ETI-related genes and nlp20-induced cell death in tobacco plants with chimeric RLP23 receptors further support this potential. The strong disease resistance conferred by RLP23 chimeras suggests that PRR engineering could approach or match the resistance levels obtained with NLRs.

In summary, our findings highlight the essential role of the IC domain in RLP-mediated immunity and its potential to enhance disease resistance across diverse plant species. RLP23 engineering achieves broad-spectrum resistance while maintaining plant growth and productivity. Unlike NLR engineering, which focuses on expanding effector recognition, RLP modifications confer robust immunity to a wide range of pathogens. RLP23’s ability to function independently of additional adaptors or helpers simplifies resistance breeding. The versatility of RLP23 across monocots, dicots, and woody plants underscores its potential for enhancing crop resilience. Future research should focus on refining RLP engineering, exploring RLP-NLR co-expression for synergistic immunity, and evaluating the long-term stability and field performance of transgenic crops to strengthen global food security.

## Methods

### Plant materials and growth conditions

*Arabidopsis thaliana* wild-type (Col-0, WT), mutant, and transgenic plants (Supplementary Table 1) were grown in soil for 7-8 weeks within a growth chamber, maintaining short-day conditions (8-hour photoperiod, 22°C temperature, and 40–60% humidity). *Solanum lycopersicum* Moneymaker wild-type and transgenic plants (Supplementary Table 1), *Nicotiana benthamiana*, rice (*Oryza sativa subsp. japonica*) and poplar (*Populus bolleana*) plants were cultivated in a greenhouse at 23°C, under long-day conditions with 16 hours of light exposure and humidity maintained at 60–70%.

### Peptides and Elicitors

Synthetic peptides nlp20, nlp24, nlp24^MbNLP1^ (GenScript), the elicitor EIX (Xylanase from *Trichoderma viride*, X3876, Sigma-Aldrich) and MoNLP1 (expressed in *Pichia pastoris)* were prepared as 10 mM stock solutions in 100 % dimethyl sulfoxide (DMSO) and diluted in water to the desired concentration before use. The GFP and Avr9 protein expressed in *Nicotiana tabacum* leaves^27^ and the recombinant SCP protein expressed in *Pichia pastoris* KM71H were purified as previously described^9^.

### Construction of RLP chimeras and mutations

IC domain swap constructs were generated using the Gibson Assembly Master Mix kit (New England Biolabs). Coding sequences for *RLP23, RLP30, Cf-9, EIX2, OsRLP1* (*LOC_Os06g04830*), and *PaRLP1* (*Potri*.*005G012100*) were cloned into pDONR207, pCR8 (Thermo Fisher Scientific) or the pLOCG vector. Using these plasmids as templates, IC domain fragments with overlapping regions were amplified and assembled with *Spe*I-digested pLOCG to produce chimeric constructs with C-terminal GFP fusions. Primers are listed in Supplementary Table 2. Putative phosphorylation sites in RLP30 (RLP30^4mut-A^) were mutated in plasmid pCR8 (containing RLP30) using overlapping PCR and primers listed in Supplementary Table 2. Epitope-tagging of non-chimeric constructs was achieved via recombination of entry clones using Gateway cloning (Thermo Fisher Scientific) into pGWB5 (C-terminal GFP-tag) or pGWB14 (C-terminal HA-tag). Constructs for NbSOBIR1-HA and SlSOBIR1-HA have been described previously^10^.

### Generation of transgenic plants

For stable integration of RLP constructs into Arabidopsis wild-type Col-0 or *rlp23*-1 mutants or tomato Moneymaker, *A. tumefaciens* GV3101 strains carrying the constructs were grown in LB medium with appropriate antibiotics. For Arabidopsis transformation, bacterial cultures were harvested, resuspended in 5 % (w/v) sucrose with 0.02 % (v/v) Silwet to an O.D. of 0.8, and inflorescences of 6–8-week-old Arabidopsis plants were dipped for 1.5 minutes^9^. 0.2 % (v/v) BASTA was used for T1 selection. For tomato transformation, bacterial cells were resuspended in 10 mM MgCl_2_ and leaf pieces were incubated in the suspension for 3 minutes before transfer to MS medium with 2 % (w/v) sucrose for 48 hours in the dark. Transgenic calli were selected on MS medium with appropriate antibiotics, then transgenic plants were moved from sterile culture to soil in a greenhouse under long-day conditions. Protein expression was confirmed by Western Blot analyses with anti-HA and anti-GFP antibodies (Thermo Fisher Scientific). To generate *NbRE02* knockout mutants, gene-specific sgRNAs (sgRNA1: 5’-CTACGGTATCCTCTCGTCAT-3’; sgRNA2: 5’-CTAGACGGGAGTTCAGGCTA-3’) were designed based on the previously reported coding sequence of NbRE02 in *N. benthamiana* using the online tool CCTop (https://cctop.cos.uni-heidelberg.de/). The pHEE401 vector carrying the gRNA sequences was introduced into *N. benthamiana* callus via *Agrobacterium*-mediated transformation. To identify CRISPR/Cas9-edited mutants, genomic DNA was extracted from mutant seedlings, and the targeted regions were verified by sequencing.

### Transient expression in plants

*A. tumefaciens* GV3101 carrying desired constructs was cultured overnight at 28°C in LB medium with selective antibiotics, then collected by centrifugation and resuspended in a solution of 10 mM MgCl_2_, 10 mM MES (pH 5.7), and 150 μM acetosyringone. The cultures were adjusted to an OD_600_ of 0.5, incubated for 2 hours at room temperature, and then infiltrated into 4-week-old *N. benthamiana* or poplar leaves. Leaf samples were collected 48 hours post-infiltration for ethylene production, ROS burst, RT-qPCR and infection assays. For transient expression in rice protoplasts, PEG-mediated transfection was performed following established protocols^28^. Briefly, 5-10 μg of plasmid DNA was mixed with 100 μl of protoplast suspension. MoNLP1 was further added to the protoplasts at a desired final concentration after 2 days.

### *In vivo* cross-linking and immunoprecipitation assays

For *in vivo* cross-linking, *N. benthamiana* leaves expressing RLP23-GFP were infiltrated with 50 nM nlp24-bio peptide, with or without 100 µM unlabeled nlp24 peptide as competitor. Five minutes post-infiltration, 2 mM EGS (ethylene glycol bis (succinimidyl succinate)) was introduced for peptide-receptor cross-linking^8^. After 20 minutes, leaf samples were harvested and immediately frozen in liquid nitrogen. For immunoprecipitation assays, constructs were transiently expressed for 48 h in *N. benthamiana* leaves. Total protein was extracted from approximately 300 mg of leaf sample and subjected to immunoprecipitation using GFP-Trap agarose beads (ChromoTek). Proteins were detected via immunoblotting with epitope-tag-specific primary antibodies (anti-GFP and anti-HA, Thermo Fisher Scientific; anti-biotin, Sigma), followed by HRP-conjugated secondary antibodies and enhanced chemiluminescence (ECL, Cytiva)^9^.

### Plant immune responses

Ethylene production was triggered in three leaf pieces floating on 0.5 ml of 20 mM MES buffer (pH 5.7) with the indicated elicitors. After 4 hours of incubation, 1 ml of air was sampled from sealed assay tubes, and ethylene levels were measured using gas chromatography (GC-14A, Shimadzu)^29^. For detection of the ROS burst, one leaf piece per well was placed in a 96-well plate (Greiner BioOne) containing 100 μl of a solution with 20 μM luminol derivative L-012 (Wako Pure Chemical Industries) and 20 ng/ml horseradish peroxidase (Applichem). Luminescence was recorded at 2-minute intervals, first for background and then for up to 60 minutes post-elicitor or mock treatment, using a Mithras LB 940 luminometer (Berthold Technologies)^9^. For RT-qPCR analysis total RNA from plant or protoplast samples was isolated using the RNeasy Plant Mini Kit (Qiagen) and used for gene expression analysis with the gene-specific primers listed in Supplementary Table 3.

### Infection assay

For bacterial infection, leaves of 6-week-old tomato plants were infiltrated with *Pseudomonas syringae* pv. tomato DC3000 (*Pst* DC3000) at a final concentration of 10^5^ CFU/ml and harvested 3 days post-inoculation as described^9^. For fungal infection, spores of *Botrytis cinerea* B05.10 were diluted to a final concentration of 10^6^ spores per ml and 5 μl drops were applied to tomato leaves. Lesion sizes were measured 2 days post-inoculation by averaging lesion diameters. Poplar leaves transiently expressing different forms of RLP23 were inoculated with a conidial suspension of *Marssonina brunnea* f. sp. *monogermtubi* (10^6^ conidia/ml)^18^. Inoculated leaves were placed on 1.5 % water agar in 9-cm Petri dishes and incubated at 25°C. Infection diameter was recorded 5 days post-inoculation. For oomycete infection, tomato leaves were inoculated with a 10 µl drop of *Phytophthora infestans* spore suspension (10^6^ spores/ml). Lesion areas were assessed 4 days post-inoculation^30^.

## Data availability

All data are available in the main text, extended data or the Supplementary Information. Source data are provided with this paper.

## Acknowledgements

We are grateful to Caterina Brancato for assistance in tomato transformation, to Matthieu Joosten for *NbSOBIR1* and *SlSOBIR1* constructs and to Rafaela Eith and Dagmar Kolb for cloning assistance. This work was supported by grants from the Deutsche Forschungsgemeinschaft (DFG) to A.A.G. (SFB766, Gu 1034/3-1), a HORIZON-MSCA-2023-PF-01-Postdoctoral Fellowship (Project: DynaCapETI) to Y.Y. and a European Research Council Starting Grant ‘R-ELEVATION’ (grant agreement: 101039824) to Y.Y. and P.D.. We also acknowledge support from the Open Access Publication Fund of the University of Tübingen.

## Author contributions

Y.Y. and A.A.G. conceived and designed the experiments; Y.Y., C.E.S. and B.L. conducted experiments; Y.Y. and A.A.G. analyzed data; L.H., J.D.G.J., D.P. and T.N. revised the manuscript and Y.Y. and A.A.G. wrote the manuscript. All authors discussed the results and commented on the manuscript.

