## Supplementary Information for "Enhancing plant broad-spectrum resistance through engineered pattern recognition receptors"

This file includes:

Supplementary Figures 1 to 4

Supplementary Tables 1 to 3

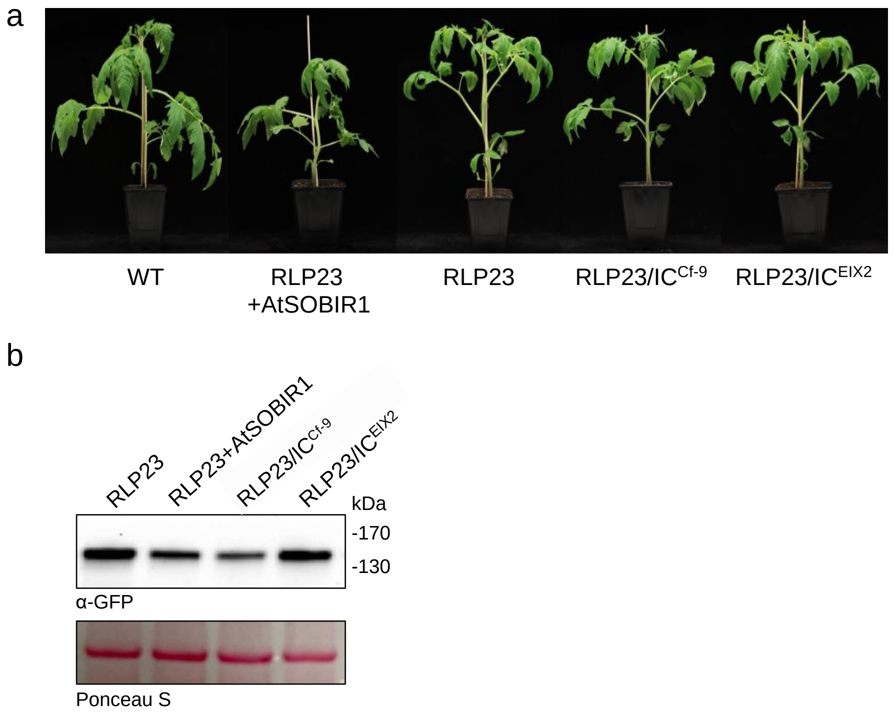


c

**RLP23** (19aa) : SYKPEWLVKIIGLNKRRKR

**Cf-9** (30aa) : STQYPAWFSRMDLKLEHIITTKMKKHKKRY

**EIX2** (37aa) : VNRSWRNAYFTFLTDMKSWLHMTSRVCFARLKGKLRN

**Figure S1: Growth, development, and morphology of transgenic tomato plants stably expressing RLP23 or RLP23 chimera are indistinguishable from wild-type plants**
**a,** Photographs of four-week-old wild-type (WT) plants or different transgenic lines stably expressing RLP23-GFP with or without AtSOBIR1-HA, RLP23/IC$^{Cf\text{-}9}$-GFP or RLP23/IC$^{EIX2}$-GFP. **b,** Western Blot analysis on tomato protein extracts from transgenic lines shown in (a) using an anti-GFP antibody, equal loading was verified by staining of the membrane with Ponceau S Red. **c,** Amino acid sequences of the intracellular domains of RLP23, Cf-9, and EIX2.

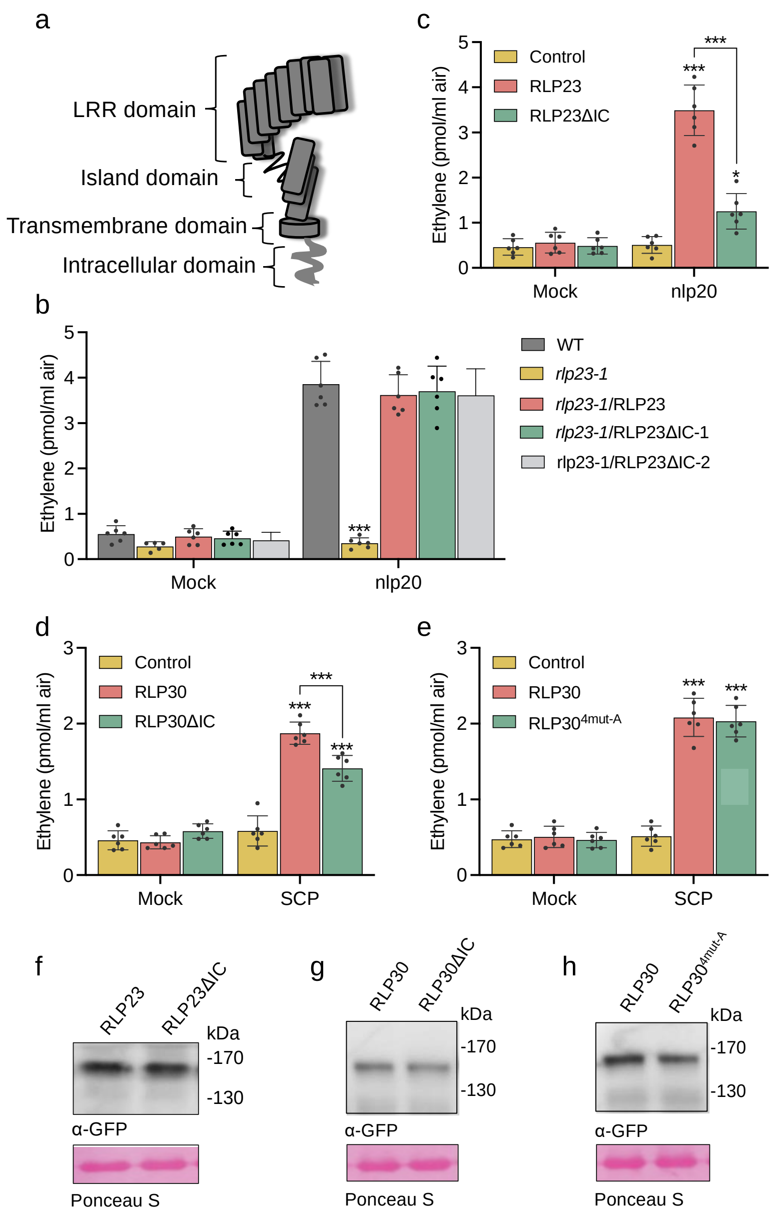


**Figure S2: Deletion of the C-terminus of RLPs doesn't affect its immune activity in Arabidopsis but reduces functionality in *N. benthamiana***

**a,** Schematic representation of the general structure of an LRR-RLP. **b,** Ethylene accumulation in wild-type (WT), *rlp23-1* and transgenic *rlp23-1* Arabidopsis plants stably expressing RLP23 or a RLP23 variant lacking the intracellular domain (RLP23ΔIC, 2 independent lines) after 4 h treatment with water (mock) or 1 µM nlp20. **c,** Ethylene accumulation in *N. benthamiana* transiently expressing GFP (Control), RLP23 or RLP23ΔIC after treatment for 4 h with water (mock) or 2 µM nlp20. **d,e,** Ethylene accumulation induced by 1 µM SCP in *RE02*-knockout *N. benthamiana* leaves transiently expressing GFP (Control), RLP30, an RLP30 variant lacking the intracellular domain (RLP30ΔIC, **d**) or a quadruple mutant in which the putative C-terminal phosphorylation sites T761, T783, T784, and S785 were replaced by alanine (4mutA, **e**). **f,g,h,** Western Blot analysis on *N. benthamiana* protein extracts from leaves shown in c-e, using an anti-GFP antibody. Equal loading was verified by staining of the membranes with Ponceau S Red. Data points are indicated as dots (*n* = 6 for b-e). Statistically significant differences from control plants are indicated (Tukey's multiple comparisons test, *P ≤ 0.05, ***P ≤ 0.001). Each experiment was repeated three times with similar results.

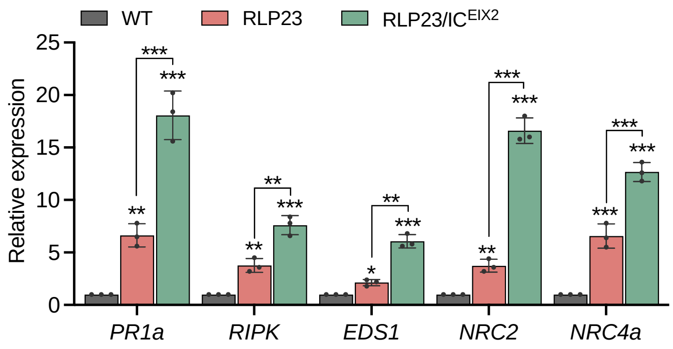


**Figure S3: Expression of RLP23/IC$^{EIX2}$ in tomato significantly enhances nlp20-induced expression of defense-related genes**

RT-qPCR analysis in wild-type (WT) and transgenic tomato plants stably expressing RLP23-GFP or RLP23/IC$^{EIX2}$-GFP after 2 µM nlp20 treatment for 24 h using gene specific primers for the indicated genes. Expression of marker genes was normalized to the levels of *SlEF1α* transcript and are presented relative to the wild-type control which was set to 1. Data points are indicated as dots (*n* = 3) and statistically significant differences from wild-type plants are indicated (Tukey's multiple comparisons test, *P ≤ 0.05, **P ≤ 0.01, ***P ≤ 0.001). Representative examples selected from three independent repetitions are shown.

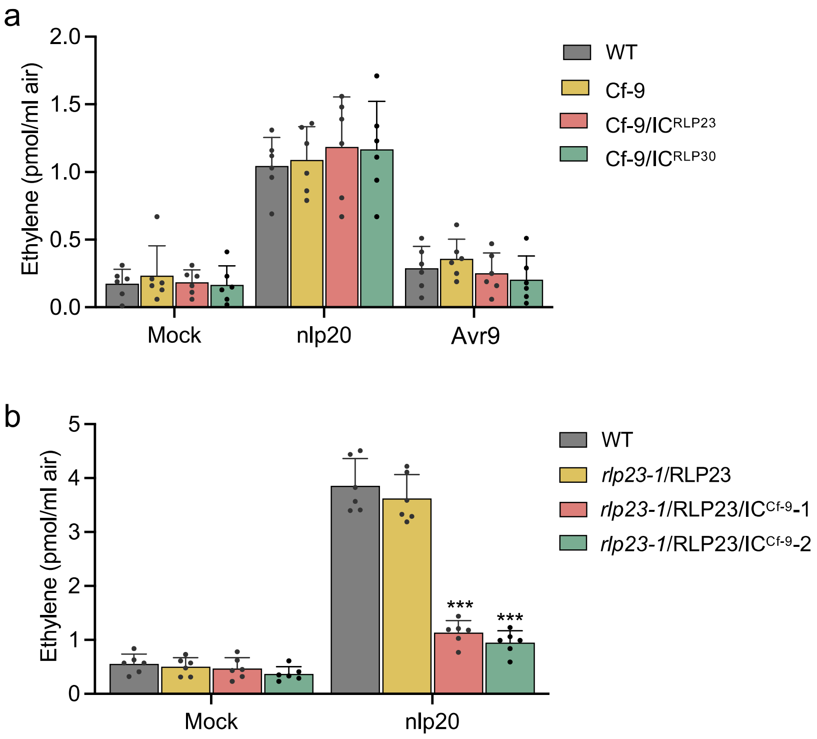


**Figure S4: Cf-9 or its chimeric versions are not functional in Arabidopsis**

**a,** Ethylene accumulation in wild-type (WT) Arabidopsis plants or transgenic lines harboring constructs for the expression of Cf-9, Cf-9/IC$^{RLP23}$ or Cf-9/IC$^{RLP30}$ after 4 h treatment with 5 µl apoplastic fluid containing GFP (Mock) or Avr9, or 1 µM nlp20. **b,** Ethylene accumulation in wild-type (WT) and transgenic *rlp23-1* lines stably expressing RLP23 or RLP23/IC$^{Cf-9}$ after 4 h treatment with water (mock) or 1 µM nlp20. Data points are indicated as dots ($n$ = 6) and statistically significant differences from wild-type plants are indicated (Tukey‘s multiple comparisons test, ***P ≤ 0.001). Each experiment was repeated three times with similar results.

**Supplementary Table 1.** Plants used in this study

| Line | Species | Description | Reference |
|---|---|---|---|
| *rlp23-1* | *Arabidopsis* | T-DNA insertion line SALK_034225 for RLP23 in Col-0 | [7] |
| *rlp23-1*/RLP23 | *Arabidopsis* | *rlp23-1* T-DNA insertion line, complemented with GFP-tagged RLP23 | [7] |
| *rlp23-1*/RLP23ΔIC | *Arabidopsis* | *rlp23-1* T-DNA insertion line, complemented with the GFP-tagged IC deletion version of RLP23 | This study |
| *rlp23-1*/RLP23/$IC^{Cf-9}$ | *Arabidopsis* | *rlp23-1* T-DNA insertion line, complemented with the GFP-tagged chimeric form of RLP23 with the Cf-9-IC domain | This study |
| Cf-9 | *Arabidopsis* | expression line of Cf-9-GFP in Col-0 wild-type | This study |
| Cf-9/$IC^{RLP23}$ | *Arabidopsis* | expression line of the GFP-tagged hybrid form of Cf-9 with RLP23-IC domain in Col-0 wild-type | This study |
| Cf-9/$IC^{RLP30}$ | *Arabidopsis* | expression line of the GFP-tagged hybrid form of Cf-9 with RLP30-IC domain in Col-0 wild-type | This study |
| *RE02* | *N. benthamiana* | Deletion line of RE02 generated via CRISPR/Cas9 | This study |
| RLP23 | Tomato | expression line of RLP23-GFP in wild-type tomato | This study |
| RLP23+AtSOBIR1 | Tomato | co-expression of RLP23-GFP and AtSOBIR1 in wild-type tomato | This study |
| RLP23/$IC^{Cf-9}$ | Tomato | expression line of the GFP-tagged RLP23 chimeric receptor with Cf-9 IC domain in wild-type tomato | This study |
| RLP23/$IC^{EIX2}$ | Tomato | expression line of the GFP-tagged RLP23 chimeric receptor with EIX2 IC domain in wild-type tomato | This study |

**Supplementary Table 2.** Primers used for cloning

| Gene | Primer name | Primer sequence (5' – 3') |
|---|---|---|
| *RLP23* | *RLP23*_F | tatggtctcatctgaacaATGTCAAAGGCGCTTTTGCA |
| | *RLP23*_R | ttggtctctccttACGCTTTCTGCGTTTATTCA |
| *RLP23*ΔIC | *RLP23*ΔIC_F | aatttactattctagtcgacctgcaggcggccgcactagtATGTCAAAGGCGCTTTTGCA |
| | *RLP23*ΔIC_R | ccccggtgaacagctcctcgcccttgctcaccatactagtAGCAATAACTTGTGCTATTG |
| *RLP23*-IC | *RLP23*-IC_F | CAATAGCACAAGTTATTGCTTCATACAAACCGGAGTGGCT |
| | *RLP23*-IC_R | ccccggtgaacagctcctcgcccttgctcaccatactagtACGCTTTCTGCGTTTATTCA |
| *RLP30*ΔIC | *RLP30*ΔIC_F | aatttactattctagtcgacctgcaggcggccgcactagtATGATTCCAAGCCAATCTAA |
| | *RLP30*ΔIC_R | ccccggtgaacagctcctcgcccttgctcaccatactagtGAAGATATGTCCAATAACTAA |
| *RLP30*-IC | *RLP30*-IC_F | AGTTATTGGACATATCTTCTTTACTGCACACAAACACGAG |
| | *RLP30*-IC_R | ccccggtgaacagctcctcgcccttgctcaccatactagtACGAGCACTTGTGGTGACTA |
| *RLP30*$^{4mut\text{-}A}$ | *RLP30*_F | ATGATTCCAAGCCAATCTAATTCC |
| | *RLP30-T761A_F* | TATCTTCTTTGCTGCACACAAACACGAG |
| | *RLP30-T761A_R* | CTCGTGTTTGTGTGCAGCAAAGAAGATA |
| | *RLP30*-Rmut-all | ACGAGCAGCTGCGGCGACTACTCT |
| *Cf-9* | *Cf-9*_F | tatggtctcatctgaacaATGGATTGTGTAAAACTTGTA |
| | *Cf-9*_R | ttggtctctccttATATCTTTTCTTGTGCTTTTTC |
| *Cf-9*-IC | *Cf-9*-IC_F | CAATAGCACAAGTTATTGCTCAACTCAATATCCAGCATGG |
| | *Cf-9*-IC_R | ccccggtgaacagctcctcgcccttgctcaccatactagtATATCTTTTCTTGTGCTTTTTC |
| *EIX2* | *EIX2*_F | tatggtctcatctgaacaATGGGCAAAAGAACTAATCCA |
| | *EIX2*_R | ttggtctctccttGTTCCTTAGCTTTCCCTTCAGT |
| *EIX2*-IC | *EIX2*-IC_F | CAATAGCACAAGTTATTGCTGTCAACCGTTCGTGGAGGAAT |
| | *EIX2*-IC_R | ccccggtgaacagctcctcgcccttgctcaccatactagtGTTCCTTAGCTTTCCCTTCAGT |
| *OsRLP1*-IC | *OsRLP1*-IC_F | CAATAGCACAAGTTATTGCTGAACTGCAAGAAATGCCTAC |
| | *OsRLP1*-IC _R | ccccggtgaacagctcctcgcccttgctcaccatactagtGTTACCTTCCGTAGTATATCCA |
| *PaRLP1*-IC | *PaRLP1*_F | CAATAGCACAAGTTATTGCTCCGCATTGGCGACGCAGGTGGT |
| | *PaRLP1*_R | ccccggtgaacagctcctcgcccttgctcaccatactagtCCTTCTGAATCTGGACAACTTG |
| *MoNLP1* | *MoNLP1*_F | CGGAATTCCTCGCGGCCCGGGACGTCATC |
| | *MoNLP1*_R | GCTCTAGACCGAAAACCTTGGCCAGGTTGTTCTG |

**Supplementary Table 3.** Primers used for RT-PCR

| Gene | Primer name | Primer sequence (5' – 3') |
|---|---|---|
| *SlEF1α* | *SlEF1α*_F | CGTGAGCGTGGTATCACCATT |
| | *SlEF1α*_R | GGTAGACCTCTCAATCATGTT |
| *SlPR1a* | *SlPR1*_F | TGGTATTAGCCATATTTCAC |
| | *SlPR1*_R | CCAGTTGCCTACAGGATC |
| *SlEDS1* | *SlEDS1*-F | GGAATTGAAGTCAGAGATGAGCTAA |
| | *SlEDS1*-R | AAAGTTCCAGCAAAAGCAAAAA |
| *SlNRC2* | *SlNRC2*-F | GACGTGGTTGGATTTGACGAAG |
| | *SlNRC2*-R | GTTACAATTCTCAGGGTGGTCTC |
| *SlNRC4a* | *SlNRC4a* -F | GCTAATCAATTTTATGCTTCCACTG |
| | *SlNRC4a* -R | CGTGATCAATGACATCGTCCTC |
| *SlRIPK* | *SlRIPK*_F | GTCATTTGACTGCAGCTAGTGA |
| | *SlRIPK*_R | GGATCTCTTGAAACCATTTTGC |
| *PaCML42* | *PaCML42*_F | CATCAATCATTGGACAGCAGTT |
| | *PaWRKY22*_R | ACCTTGAATGCCTCTGACAAAT |
| *PaWRKY22* | *PaCML42*_F | GTCTAAAGGGATGTTTGGCAAG |
| | *PaWRKY22*_R | TTTAGGCATAGATGGCTTGGTT |
| *PaRAP2.6* | *PaRAP2.6*_F | ACTGTGCTCATGGTGATTTGTC |
| | *PaRAP2.6*_R | GCTCTTCTCTGCAGGTTTCATT |
| *PaPR1* | *PaPR1*_F | TCAATGCCCACAATAATGCTCG |
| | *PaPR1*_R | TAAGATCACCACTACCTCCTGC |
| *PaEF1α* | *PaEF1α*_F | CCTGGACATCGTGACTTTATCA |
| | *PaEF1α*_R | GTCCATCTTGTTACAGCAGCAG |
| *OsWRKY70* | *OsWRKY70*_F | CCGCTGCTGTTTTGATCATCT |
| | *OsWRKY70*_R | CCGCTGCTGTTTTGATCATCT |
| *OsPAL* | *OsPAL*_F | CTACCCGCTGATGAAGAAGC |
| | *OsPAL*_R | GAACCTTGTTCAGCTCCTCG |
| *OsPR1a* | *OsPR1a*_F | TCGTATGCTATGCTACGTGTTT |
| | *OsPR1a*_R | CACTAAGCAAATACGGCTGACA |
| *OsMAPK6* | *OsMAPK6*_F | CGCACGCTCAGGGAGATC |
| | *OsMAPK6*_R | GGTATGATATCCCTTATGGCAACAA |
| *OsActin* | *OsActin*_F | TTATGGTTGGGATGGGACA |
| | *OsActin*_R | AGCACGGCTTGAATAGCG |
